# Developmental Ramifications of the Screen Age: Associations between Screen Time and Neural Maturation in a Massive Adolescent Cohort

**DOI:** 10.1101/2023.09.27.559772

**Authors:** Alexander S. Atalay, Benjamin T. Newman, T. Jason Druzgal

## Abstract

A growing body of literature associates increases in electronic screen time with a vast array of psychological consequences amongst adolescents, but little is known about the neurological underpinnings of this relationship. This longitudinal study examines structural and diffusion brain MRI scans from the Adolescent Brain Cognitive Development (ABCD)Study: a large multi-site study with thousands of participants. By assessing both gray matter density (GMD) and grey matter measurements of diffusion microstructure in the adolescent brain, we describe how the developmental trajectory of the brain changes with screen-based media consumption at the sub-cellular level. Grey matter microstructure was measured across 13 bilateral regions functionally implicated with screen time use, and associated with either the control or reward system. After controlling for age, sex, total brain volume, scanning site, sibling relationships, physical activity, and socioeconomic status, this study finds significant positive correlations between increased screen time and axonal signal across 6 of the 13 regions while also finding significantly decreased intracellular signal in 8 regions. Comparing these associations to normal developmental trajectories suggests adolescent age-related brain development may be accelerated by increased screen time in brain areas associated with reward processing while age-related brain development may be decelerated in regions of the control system. Highlighting the sensitivity of microstructural analysis, no significant cross-sectional or longitudinal relationship with increased screen time was found using GMD, or fractional anisotropy. This work suggests that increased screen usage during adolescent development has a complex association with brain tissue that cannot be completely described by traditional quantifications of tissue microstructure.

## Introduction

Adolescence is a crucial period for the growth and maturation of the brain and the development of cognitive capabilities that will affect the remainder of the lifespan. As a result of technological advances and the COVID-19 pandemic, screen time usage is at an all-time high.^1^ In adolescents, increased screen time has been linked to lower psychological well-being^2^, but little is known about the relationship between increased screen time and neural development, and previous neuroimaging studies on the subject have focused on either small or cross-sectional subject pools.^3^ This study examines structural and diffusion MRI scans from the longitudinal Adolescent Brain Cognitive Development (ABCD) study cohort to describe the effects of screen time on both grey matter density (GMD) and diffusion MRI (dMRI) tissue microstructure in the adolescent brain. 13 regions of interest (ROIs) identified by previous studies were selected for investigation based on either their functional involvement during screen time usage or structural associations with isolated screen-oriented activities in adolescence.^3^ These ROIs were classified based on primary involvement in either the cognitive control system or reward processing system, two networks that have demonstrated substantial relationships with screen-based activates in adolescents across multiple studies.^3^

### The Effects of Screen Time on Adolescent Development

The adolescent brain undergoes substantial neuroanatomical changes at both global and cellular scales during development. Developing brains typically present increases in total brain volume accompanied by non-linear global increases in white matter (WM) volume and decreases in the volume of gray matter (GM).^4–6^ While these relationships are typically spatially and temporally dependent on region and age, they offer insight into the functional reorganization of the brain during development, and non-traditional development trajectories can predict a variety of neuropsychiatric pathologies such as schizophrenia and bipolar disorder.^7^ Accompanying the growth of cognitive abilities, the cognitive-control system, responsible for the ability to select behaviors based on thoughts, emotions, and social context,^8^ matures substantially during adolescence. Furthermore, the maturation of affective control–the application of cognitive control to emotional contexts–drives developmental changes in emotional regulation tendencies during adolescence, specifically relying on changes in connectivity between prefrontal regions and regions implicated in emotion and reward processing.^9,3^

Increased screen time has been associated with a host of negative psychological and physiological outcomes in adolescents, including poor sleep, high blood pressure, obesity, poor stress regulation, and insulin resistance.^7^ Symptoms of attention deficit/hyperactivity disorder (ADHD), a neurodevelopmental disorder characterized by inattention, hyperactivity, and impulsivity, have also been linked to screen media consumption.^10^ Relationships have been found between overall screen time and increased symptoms of depression and suicidal behavior among adolescents.^11^ While these studies demonstrate a clear association between screen time and adverse health outcomes, the mechanism by which screen time influences brain development is largely undescribed. Despite the growing concern surrounding general screen use in adolescents and the mounting evidence for its consequences, little is known about the neurological basis for screen driven changes during development.

Neuroimaging research focused on structural changes associated with screen time has been heterogeneous, with significant associations primarily being found when subjects are followed longitudinally. In a cross-sectional analysis of frequency of internet use and development of brain structures in adolescents, there were no significant associations between a higher frequency of internet use and regional grey matter or white matter volumes calculated through voxel-based morphometry (VBM). However, longitudinally, a higher frequency of internet usage predicted change in regional grey matter and white matter volumes in a widespread anatomical cluster, including the orbitofrontal cortex (OFC), temporoparietal junction (TPJ), insula, and amygdala.^12^ Cross-sectional research focused on the effects of screen time on white matter microstructure quantified through fractional anisotropy (FA) found no significant relationships.^13^ Previous research has also evaluated the relationship between screen time and functional connectivity. In a 2018 study involving 19 10-year-old children, screen time was related to lower connectivity between the left visual word form area and the anterior cingulate cortex (ACC), inferior frontal gyrus (IFG), and insula.^14^ Research investigating the effects of media-related tasks on the adolescent brain using a task-based fMRI approach have established robust relationships between social media use and reward network recruitment^15,16^, as well as the association between sedentary screen time with decreased impulse-control.^17^ In research comparing adolescent subjects with internet addictions to healthy controls, relationships were found between problematic internet usage and lower gray matter density in the left ACC, posterior cingulate cortex, and insula^18^, as well as a reduction in cortical thickness in the OFC.^19^ Finally, in a network-scale analysis, frequency and duration of screen-based media consumption was correlated with lower overall efficiency of the cognitive control system.^3^

A previous review of brain MRI studies in the developmental screen time literature identified 13 bilateral ROIs demonstrating longitudinal relationships with increased screen time based on associations with either task-related or task-unrelated media usage.^3^ Of the 13 regions of interest analyzed, 7 are regions involved in the cognitive control system, and 6 are involved in the reward system. The control system regions analyzed in this study include the ventromedial prefrontal cortex (vmPFC), orbitofrontal cortex (OFC), inferior frontal gyrus (IFG), superior parietal lobule (SPL), inferior parietal lobe (IPL), temporal-parietal-junction (TPJ), and the anterior cingulate cortex (ACC). The reward system ROIs are located in the insula, dorsal ACC (dACC), amygdala, caudate nucleus (CAU), putamen, and nucleus accumbens (NAc). A summary of the regions of interest, their functional associations, and their neuroanatomical arrangement, can be found in Fig. 1 and Table 1.

**Table 1:** Regions of interest selected for this study based on previous functional or structural associations with screen-media consumption. Region indices refer to the labels used in Fig 1. Regions are divided based on their roles in either control or reward systems.^3^

| Figure 1 Index | Region | Associated System |
| --- | --- | --- |
| 1 | vmPFC | Control |
| 2 | OFC | Control |
| 3 | IFG | Control |
| 4 | SPL | Control |
| 5 | IPL | Control |
| 6 | TPJ | Control |
| 7 | ACC | Control |
| 8 | Insula | Reward |
| 9 | dACC | Reward |
| 10 | Amygdala | Reward |
| 11 | CAU | Reward |
| 12 | Putamen | Reward |
| 13 | NAc | Reward |

**Figure 1:**
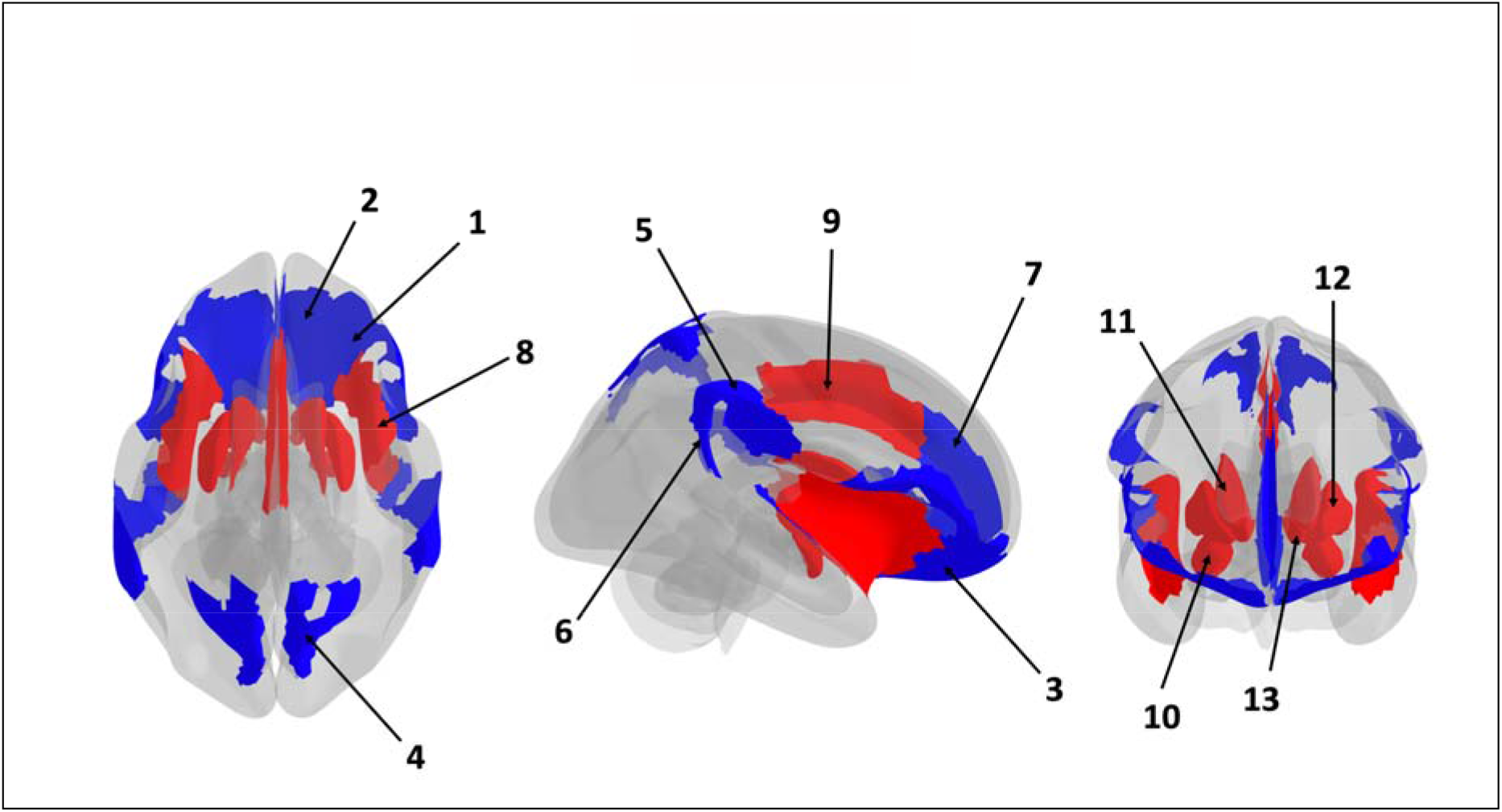
ROI map overlay showing the spatial relationships between selected regions. ROIs are color filled based on associations with control (blue) or reward (red) systems.

### Microstructural Analysis

Modern MRI analysis pipelines allow for quantification of detailed neuroanatomical characteristics through structural imaging. Two such techniques are Voxel-Based Morphometry^20^ (VBM), which allows for the calculation of cortical or subcortical gray matter density (GMD) from structural MRI, and 3-Tissue Constrained Spherical Deconvolution (3T-CSD)^21,22^, which can quantify sub-cellular components of brain microstructure from dMRI. GMD measures the signal strength in grey matter regions of the brain, acting as a quantification for the general tissue environment. 3T-CSD provides even deeper insight, estimating the proportion of three cellular microenvironments found in the brain: intracellular isotropic (ICI), intracellular anisotropic (ICA), and extracellular isotropic (ECI). These metrics represent signal fractions of water inside cell bodies and structures, water inside axons, and freely diffusing extracellular water, respectively. By tracking the changes in these tissue environments, we can gain insight into what is happening in the brain at a subcellular level.

Based on previous results and established typical developmental trajectories, we hypothesize that subjects with increased screen time will demonstrate reduced or delayed neural development longitudinally. More specifically, we expect to observe a significantly smaller decrease in GMD and ICI signal fraction as well as a significantly smaller increase in ICA signal fraction than observed during typical development.

## Methods

### Participants

This study utilized data obtained from the Adolescent Brain Cognitive Development (ABCD) study^23^, a publicly available, longitudinal, neuroimaging and demographic study examining child development and factors leading to substance abuse. Due to storage and computational limitations and to avoid manufacturer and sequence differences, including field gradient strength, TE, and TR, that have previously been demonstrated to affect outcome 3 Tissue, Constrained Spherical Deconvolution model (3T-CSD) signal fraction results, only subjects from the most common scanner type, Siemens, proceeded to analysis.^24^ Subjects missing any relevant data field were removed from the analysis.

Following a semi-automated quality control process including removing subjects without completed diffusion or T1-weighted scans, excessive motion, other imaging artifacts, and successful registration into a cohort-specific template space 3284 and 3952 subjects remained for final analysis from baseline and follow-up timepoints respectively. This includes a longitudinal crossover of 2666 subjects, and a total 4554 unique subjects. There were significantly more males than females in both baseline and follow-up samples (1741 males (53.2%) versus 1527 females (46.8%), p<0.001 at baseline; 2122 males (53.7%) versus 1814 females (46.3%), p<0.001 at follow-up). The age ranges at baseline were the same for the female cohort and the male cohort (107.0 to 132 months), and there was no significant difference between the average age of females (119.5 months ± 7.4 SD) and the average age of males (119.9 months ± 7.5 SD) (p=0.086). While the age ranges were slightly different at the follow-up timepoint (129 to 162 months for females vs 127 to 160 months for males), there was no significant difference between the average age of females (142.8 months ± 7.6 SD) and the average age of males (143.2 months ± 7.7 SD) (p=0.080) in the follow-up sample.

### Demographic Data

Screen time data was acquired through the ABCD Youth Screen Time Survey (STQ): a parent-reported questionnaire administered at both the baseline and two-year follow up timepoints. Respondents were asked to indicate the number of hours per day their child participated in an array of screen-based activity such as watching TV shows or movies, watching videos (such as YouTube), playing video games, texting, visiting social networking sites, and video chatting. Reported average screen usage during weekdays and weekend days across all activates were aggregated with a weighted average to estimate the total hours per week of screen time. The screen time usage between baseline and follow up timepoints was then averaged for each subject and used as a predictor variable for linear regression models. Exact survey questions were changed between baseline and follow up timepoints to account for an increased granularity in screen time usage, but aggregating methods were fundamentally the same across timepoints.

**Figure 2:**
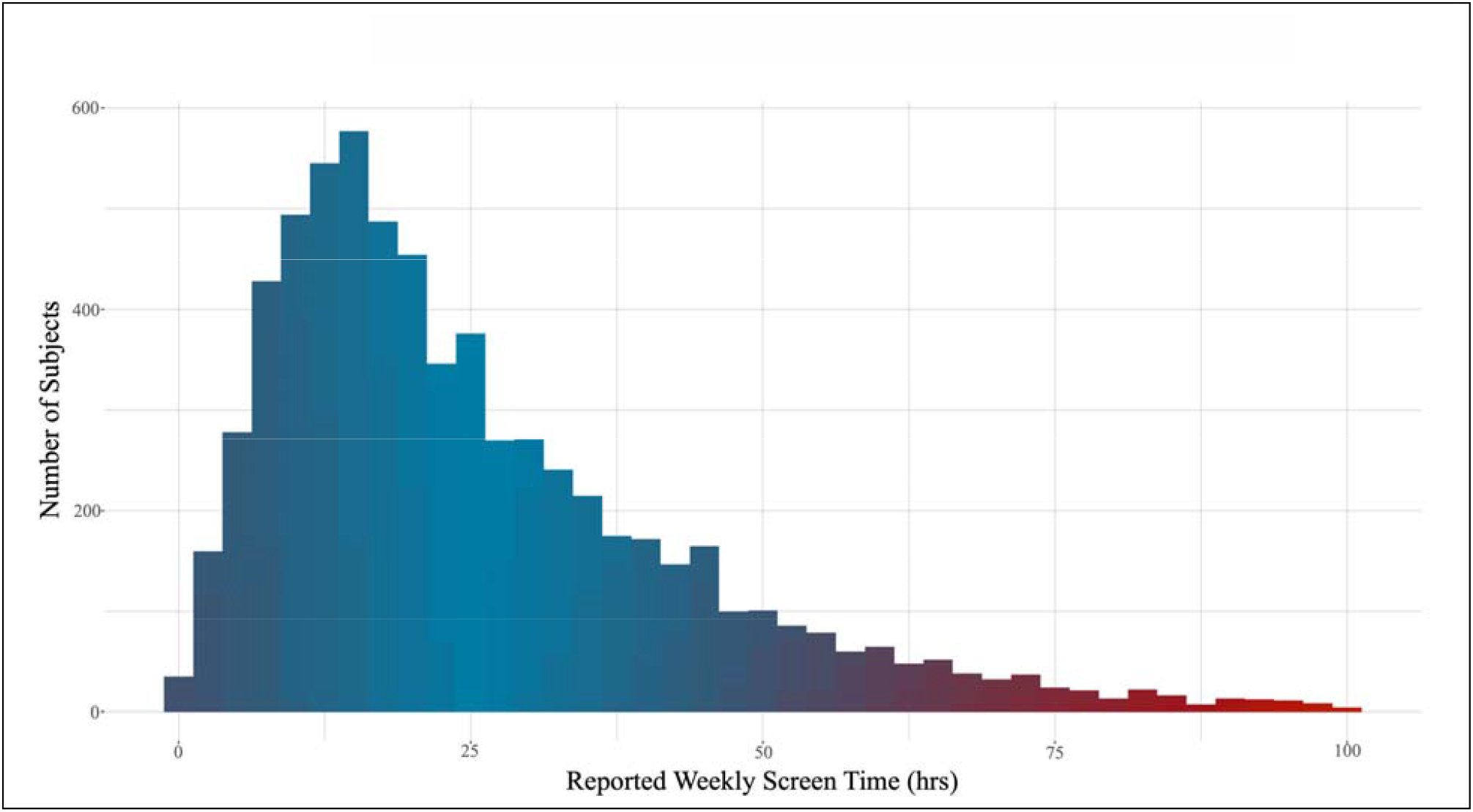
Average Weekly Screen Time Histogram. A histogram of average weekly screentime hours by subjects included in this study. Average weekly screentime hours were calculated by summation of parent responses to weekday and weekend screen time uses by activity and averaged across timepoints. The average reported weekly screen time across all subjects is 25.6 hours.

Sports involvement data, used as a proxy for physical activity, was acquired through the ABCD Longitudinal Parent Sports and Activities Involvement Questionnaire (SAIQ): a parent-reported questionnaire administered at both the baseline and two-year follow up timepoints.

Respondents were asked to indicate the number of hours per day their child participated in a wide variety of physical activates and sports such as soccer, surfing, and horseback riding. Reported weekly hours for each activity was summed for each subject and used as a control for linear regression models.

Household income data, used as a proxy for socioeconomic status, was acquired through the ABCD Parent Demographics Survey, in which parents report total combined household income for the past 12 months. This survey was administered at both baseline and follow up timepoints. The ABCD Study includes an oversampling of families with twins and the final cohort included 1070 twins, 19 triplets (drawn from 7 families of triplets, but in two families one subject did not pass quality control) and 563 non-twin siblings^25^.

### Imaging Data

Unprocessed T1-weighted ‘Magnetization Prepared Rapid Acquisition Gradient Echo’ (MPRAGE) MRI pulse sequences were obtained from the ABCD study. These images were acquired with an isotropic voxel size 1.0 × 1.0 × 1.0 mm^3^ with TE = 2.88 and TR = 2500 with a flip angle of 8 degrees and an field of view of 256 × 256 mm^2^.^23^

As described in Newman et al., 2020^26^: Unprocessed dMRI images were obtained from the ABCD study and were acquired with a multiband accelerated sequence that had an isotropic voxel size 1.7 × 1.7 × 1.7 mm^3^ with TE = 88 ms and TR = 4100 ms. Using a multi-shell protocol 7 images were acquired at b= 0, 6 directions were acquired at b=500 s/mm^2^, 15 directions were acquired at both b=1000 s/mm^2^ and at b=2000 s/mm^2^, and 60 directions were acquired at b=3000 s/mm^2^.^23^ Two bidirectional field maps with reverse phase encoding were obtained at b=0 with identical isotropic voxel sizes, TE, and TR for use in distortion correction.

Total brain volume and fractional anisotropy (FA) data were obtained from the ABCD study baseline and follow-up releases. Volumetric data was calculated using an automated processing pipeline in Freesurfer version 6.0.1.^27^ FA data was calculated through a diffusion tensor model using a standard, linear estimation approach with log-transformed diffusion-weighted signals as described in Palmer et al, 2022.^28^

### Regions of Interest

ROIs were selected based on previous association with screen time usage in functional and structural neuroimaging studies as described by Marciano et al 2021.^3^ Primary clusters of activation identified by these previous studies were localized to bilateral ROIs available in the Destrieux cortical atlas.^29^

### Imaging Preprocessing and Analysis I: GMD from T1W MRI

VBM analysis was performed identically to Barrett et al., 2018^30^: Using acquired magnetization-prepared rapid gradient-echo T1 sequence (MPRAGE), we applied voxel-based morphometry methodology.^20,31^ Briefly, images were spatially normalized to standard stereotactic space through both an affine and high-dimensional nonlinear registration. MRI scans were segmented into gray and white matter and high-dimensionally fit to the Montreal Neurological Institute (MNI) standard space with the CAT12 toolbox (dbm.neuro.uni-jena.de/cat/) in conjunction with SPM12 (fil.ion.ucl.ac.uk/spm/software/spm12/) in MATLAB (MathWorks, Natwick, MA). To preserve absolute volume of gray matter, segmented images were multiplied by the relative voxel volumes contained within the jacobian determinant matrix of the deformation field. Regional GMD was measured as an average of each Destrieux atlas map within a ROI.

### Imaging Preprocessing and Analysis II: Tissue Microstructure from dMRI

3T-CSD analysis was performed identically to Newman et al., 2022^32^: Image preprocessing was performed consistent with prior protocols that have been shown to result in consistent and reliable signal fraction measurements.^24^ All dMRI images were corrected for thermal noise using the “dwidenoise” command implemented in MRtrix3.^33^ Gibbs rings were then removed with the “dwidegibbs” MRtrix3 function. The FSL package (“topup” and “eddy”) was subsequently applied to correct for susceptibility-induced (EPI) distortions, eddy currents, and subject motion, including the Gaussian replacement of outliers^34–37^. Finally the preprocessed images were upsampled using MRtrix3 to alter the resolution to 1.3 × 1.3 × 1.3 mm^3^ isotropic voxel size, in order to bring their spatial resolution closer to the high-resolution acquisition of images in the Human Connectome Project which serves as a benchmark for high quality diffusion data^38,39^. The b=0 and b=3000 s/mm^2^ shells were then extracted to form a single-shell image set suitable for single-shell, 3 tissue constrained spherical deconvolution (SS3T-CSD). This step was performed because prior investigations have suggested SS3T-CSD is superior at differentiating between brain regions compared to multi-shell multi-tissue CSD at b=3000 s/mm^2 24^.

Brain masks were obtained for all subjects by performing a recursive application of the Brain Extraction Tool^40^. Response functions from each of the three tissue types were estimated from a randomly selected subset of nearly 500 subjects and averaged to produce a single set of tissue response functions^41^. The response functions were selected via an unsupervised method described by Dhollander et al., (2016) briefly, the WM tissue response function was selected from an FA thresholded mask, the CSF tissue response function was selected in voxels with the highest signal decay metric between the averaged b-0 and b=3000 s/mm^2^ shells, and the GM tissue response function was selected from voxels closest to the median voxel-wise signal decay metric, after a conservative GM mask was constructed. SS3T-CSD was then performed using the average response functions with MRtrix3Tissue, a fork of MRtrix3, to estimate an anisotropic WM-like (represented by a complete WM fiber orientation distribution (FOD), i.e. all fit signal from WM fibers); an isotropic, intracellular GM-like (the signal that is fit by neither WM or CSF response functions, and more closely fit by an isotropic but less b-value dependent signal response function); and an isotropic, extracellular CSF-like compartments^24,^. SS3T-CSD is functionally a specialized optimizer that iteratively performs a linear least squares fit of the response functions similarly to multi-shell multi-tissue CSD (MSMT-CSD). Initially, the isotropic CSF and GM response functions are fit to the diffusion signal, with WM as a prior constraint. This calculation yields an underestimate of CSF from the CSD fit. The next step fits anisotropic WM and isotropic GM response functions with the previously calculated CSF as a constraint, this also yields an underestimate of WM from the CSD fit. Over continuous iterations, this method attempts to assign signal repeatedly to either WM or CSF compartments from the GM compartment, repeatedly performing a cleaner separation as signal is assigned and remaining signal is run through the algorithm again. An example of this general pipeline, including preprocessing steps, is available at https://3tissue.github.io/doc/single-subject.html. Each subject’s three tissue compartments were then normalized to sum to 1 on a voxel-wise basis, resulting in the final three-tissue signal fraction maps. Summing the spherical harmonic coefficients on a rotational-invariant voxel-wise basis provides the added benefit of harmonizing inter-scanner and inter-subject signal intensity differences while preserving between-subject biological variation^43^.

A cohort specific template was constructed from a sex-balanced random selection of 40 subjects’ WM-FODs at both baseline and the followup timepoint using symmetric diffeomorphic registration of the FOD themselves and implemented in MRtrix3 with the “population_template” function. Each subject was then individually registered to the cohort template using an affine, followed by a nonlinear registration guided by the WM FODs themselves using an apodized point function^45^. The resulting warp was used to move each of the subjects’ three-tissue signal fraction maps into the common template space. The cohort template was then registered to stereotaxic space with a similar FOD-based diffeomorphic registration procedure to a b-value matched version of the NTU-DSI-122 template to establish registration with stereotaxic MNI-space.^26^ ROIs were then warped into the common template space and mean 3T-CSD values were measured within each ROI. For FA the relevant regions from the Destrieux atlas provided by the ABCD study were averaged together to create the equivalent of the fMRI-defined ROIs by weighting each individual constituent ROI’s mean FA using that region’s volume.

## Statistical Analysis

### Data Summarization

Categorical variables are summarized by frequencies (n) and percentages (%). Continuous scaled variables are summarized by the mean, standard deviation, and range of the empirical distribution.

### GMD, FA, and Tissue Signal Fraction Regression Analysis

Type II ANOVA models were used to predict mean ROI GMD, FA, and tissue signal fractions (i.e. extracellular isotropic; or intracellular isotropic; or intracellular anisotropic) as a function of a subject’s average weekly screen time (hrs/week), age (years), sex (female, male), total brain volume (TBV, cm^3^), combined family income, and physical activity (hrs/week).

In all models, each child’s family and sibling relationship was accounted for as a nested random effect within site to ensure twins, triplets, and siblings were not biasing results following recommendations by Saragosa-Harris et al., (2022). All of the model predictor variables and interactions that were selected a priori based on scientific merit, and were identical between each imaging derived metric of interest, displayed as equation 1:

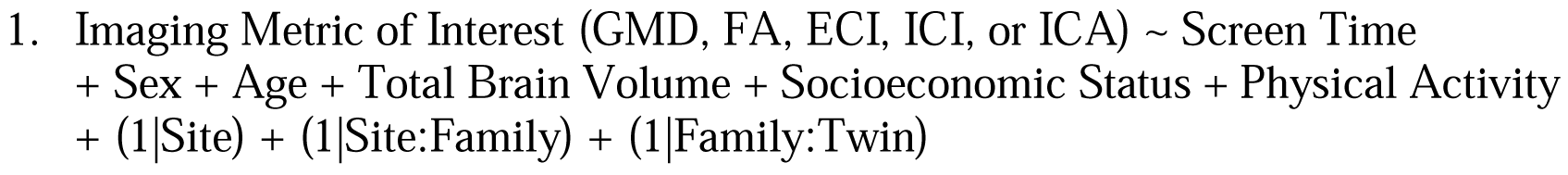

For each ROI, the concomitant variable adjusted association between tissue signal fraction and average ST was quantified by the regression slope coefficient estimate associated with weekly ST. For each ROI, a null hypothesis test was performed to test the null hypothesis that the slope of the association between the tissue signal fraction and weekly ST is equal to 0, versus the alternative that the slope of the association between the tissue signal fraction and screen time is not equal to 0. The complete set of p-values from the 13 different ROI were then subjected to the Holm-Bonferroni method for multiple comparisons to identify those ROI, in which the ANOVA type II F-test p-value of the null hypothesis test was less than the adjusted p-value for each hypothesis. This correction was performed for each imaging metric separately.

## Results

### GMD and FA Regression Analyses

The ANOVA summaries for the mixed-effect regression models that were used to account for random nested effects from sibling relationships and random effects from scanner site predict GMD and FA as a function of the subject’s average weekly screen time, and the subject’s age, sex, household income, physical activity, and total brain volume are summarized in Table 3: displaying the number of ROIs that had a significant association with each of the predictor variables. While are no significance relationships between subject weekly screen time and GMD in any of the 13 ROIs analyzed, age, sex, total brain volume, and household income are significant predictors across multiple regions. Similarly, while only the ventromedial prefrontal cortex showed significant associations between screen time and FA, age, sex, physical activity, total brain volume, and household income are significant predictors across multiple regions.

**Table 3:**
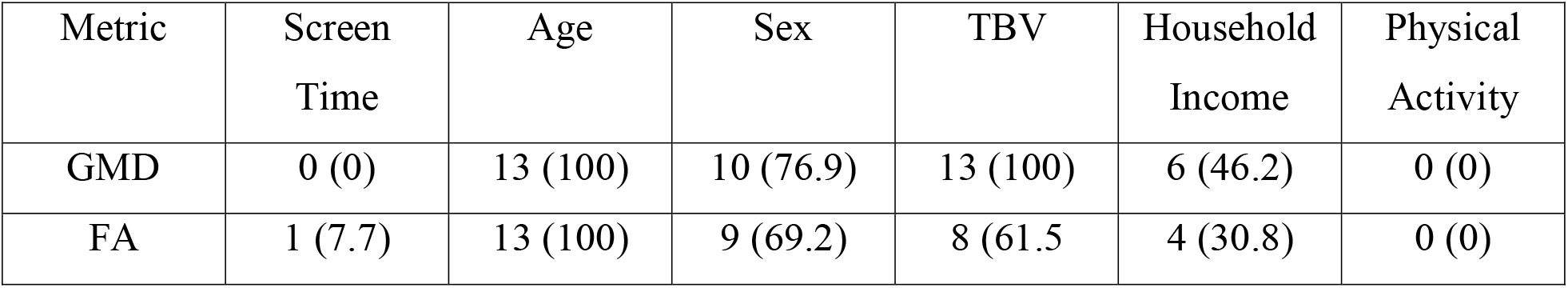
Number of ROI (out of 13 total) where each associated variable remained significantly associated with GMD and FA metrics after implementation of the Holm-Bonferroni method for multiple comparisons (p<0.05).

Longitudinal relationships between both GMD and FA data and age are observed across all 13 ROIs. Subject age is inversely related to the GMD of every ROI with the exception of the Amygdala. Similarly, subject age is inversely related with the FA of every ROI analyzed.

### Tissue Signal Fraction Regression Analyses

The ANOVA summaries for the mixed-effect regression models that were used to account for random nested effects from sibling relationships and random effects from scanner site predict tissue signal fraction (i.e. extracellular isotropic CSF-like; or intracellular isotropic GM-like or intracellular anisotropic WM-like) as a function of the subject’s average weekly screen time, and the subject’s age, sex, household income, physical activity, and total brain volume are summarized in Table 4: displaying the number of ROIs that had a significant association with each of the predictor variables.

**Table 4:** Number of ROI (out of 13 total) where each associated variable remained significantly associated with each tissue signal fraction compartment after implementation of the Holm-Bonferroni method for multiple comparisons (p<0.05).

| Signal Fraction | Screen Time<br># (%) | Age<br># (%) | Sex<br># (%) | TBV<br># (%) | Family Income<br># (%) | Physical Activity<br># (%) |
| --- | --- | --- | --- | --- | --- | --- |
| ECI | 1 (7.7) | 12 (92.3) | 4 (30.8) | 5 (38.5) | 7 (53.8) | 3 (23.1) |
| ICI | 8 (61.5) | 12 (92.3) | 1 (7.7) | 8 (61.5) | 7 (53.8) | 0 (0) |
| ICA | 6 (46.2) | 13 (100) | 3 (23.1) | 4 (30.8) | 4 (30.8) | 0 (0) |

Significant longitudinal associations between screen time and 3T-CSD tissue microstructure were found in 8 regions. A significant positive correlation was observed between screen time and ICA signal fraction in the vmPFC, the orbitofrontal cortex, the SPL, the TPJ, the caudate nucleus, and the putamen. In these regions and the IFG and IPL, a correspondingly significant negative correlation between screen time and ICI signal fraction was observed.

Longitudinal relationships between 3T-CSD metrics and age are widespread and show mixed congruency with the relationship between 3T-CSD metrics and screen time. Subject age is positively correlated with ICA signal across all 13 ROIs and ICI signal in the vmPFC, the IFG, the SPL, IPL, and TPJ. Age is negatively correlated with ICI signal in the ACC, insula, dACC, amygdala, CAU, putamen, and NAc. Congruency between the associations of screen time on tissue microstructure and age on tissue microstructure is defined as whether or not the slopes of these relationships share the same direction (positive or negative). Congruency is observed in the ICA signal fraction of every significant region as well as ICI signal fraction in the putamen and CAU. Incongruency is observed in the ICI signal fraction of the vmPFC, IFG, SPL, IPl, and TPJ. Age, sex, total brain volume, and household income were also shown to significantly contribute to tissue microstructure.

**Figure 3:**
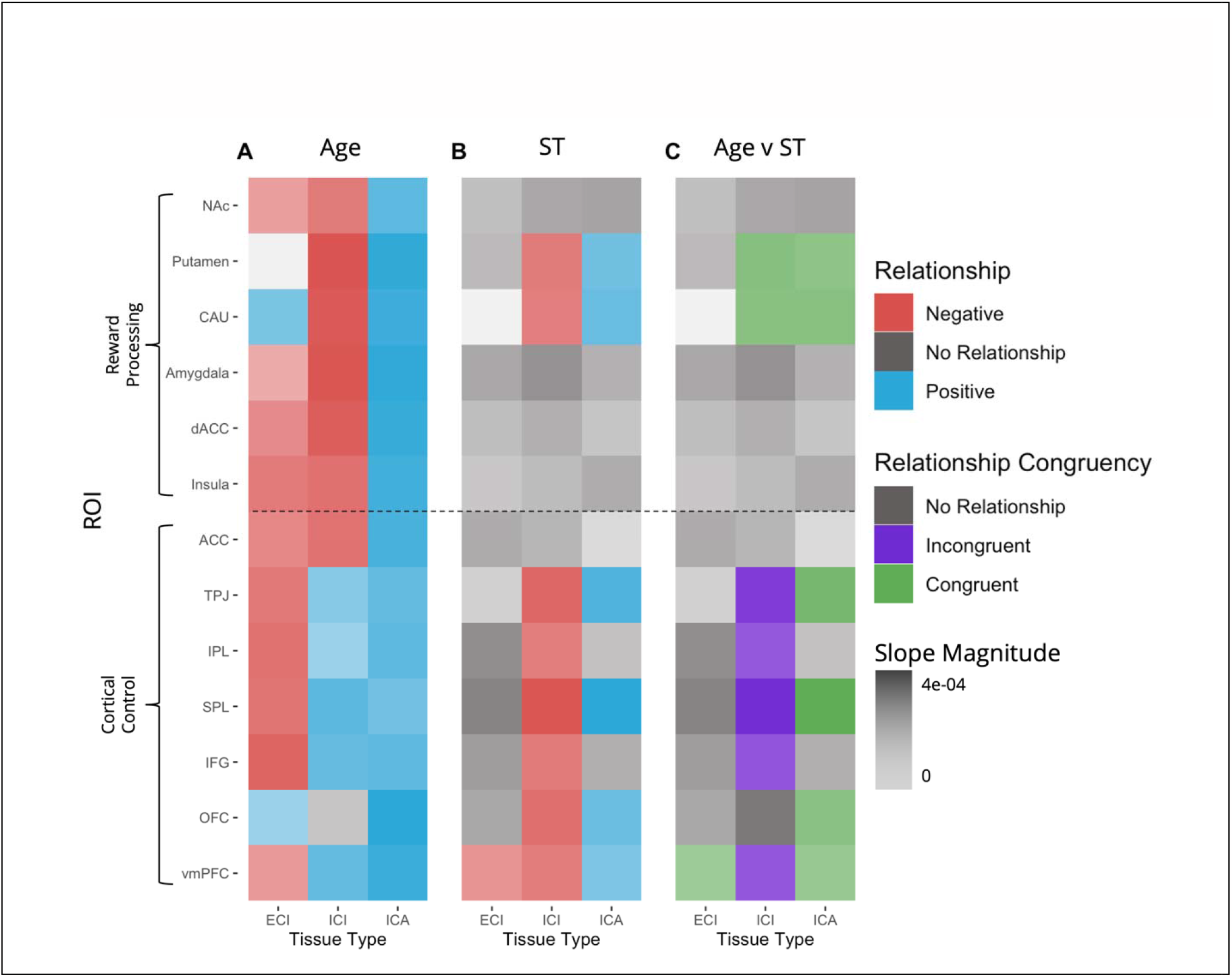
Relationships between Age and Screen Time on Regional Tissue Microstructure. Regional relationships of both age and screen time (ST) on tissue microstructure showcasing relationship direction and strength – organized by functional associations. **A)** Regional relationships between age and tissue microstructure characterizing typical development. All but 2 region/signal fraction combinations demonstrate significant relationships with age that vary in direction and strength. **B)** Regional relationships between average weekly screen time and tissue microstructure. 8 out of 13 ROIs demonstrate significant relationships with screen time across at least one signal fraction. **C)** Regional comparisons between age and screen time on microstructure. Congruency is defined as whether the relationship of age and of screen time on microstructure are in the same direction (both positive or both negative) or different directions (one positive and one negative) for a given region and tissue type.

**Figure 4:**
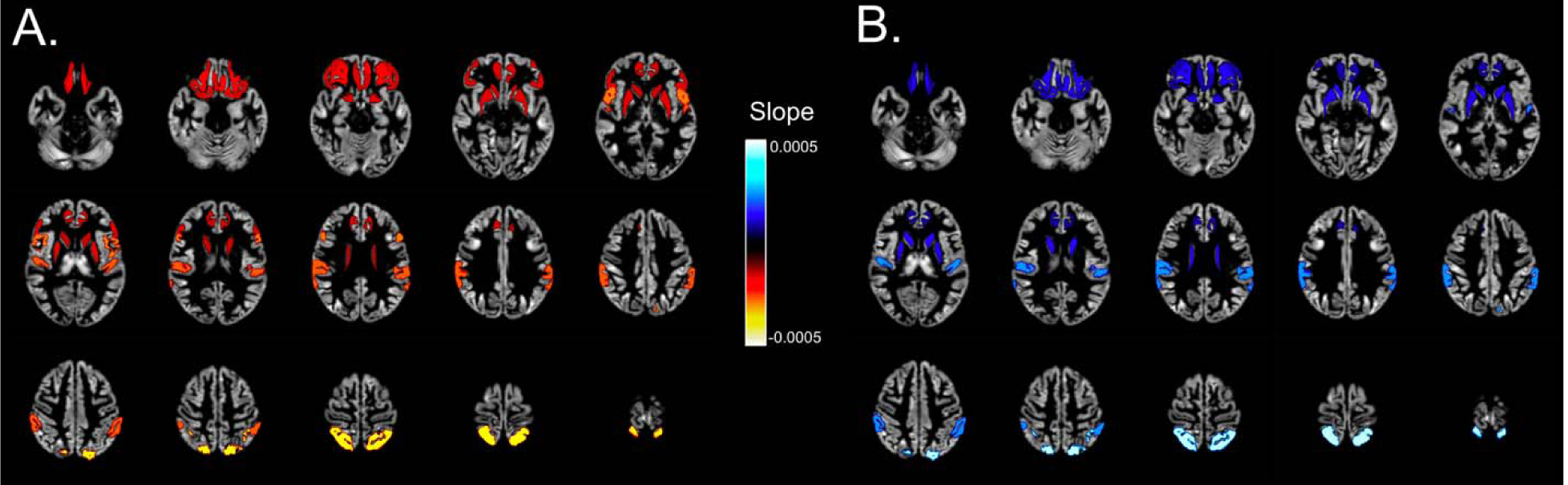
Significant Regional Relationships between Screen Time and Microstructure. Display of significant adjusted intracellular isotropic (**A**) and intracellular anisotropic (**B**) signal fraction model slopes from analyzed ROIs. ROI maps are colored by slope (representing the change in signal fraction per hour of weekly screen usage) and displayed on the cortical ribbon of the cohort specific template.

## Discussion

In this longitudinal MRI study of 2666 adolescents from the ABCD study we have identified a relationship between parent reported screen media usage and several measurements of brain tissue microstructure across brain regions involved in the control and reward systems.

Our observations about the effects of age on typical neural development on tissue microstructure in adolescents agrees with previous findings and paints an important contrast to our results on the effects of increased screen time. In all but two regions, we found significant relationships between subject age and tissue microstructure across tissue types. Increased subject age has a negative relationship with ECI signal fraction in every region except for the putamen (no relationship), the CAU (positive relationship) and the OFC (positive relationship). This suggests that as adolescents experience typical development, the amount of CSF/free water in these regions tends to decrease with age. Age has a consistent positive relationship with the ICA signal fraction in all ROIs, suggesting that, in these regions, the proportion of myelinated fibers increases with age. Finally, subject age has a region-dependent effect on the ICI signal fraction. In every control region analyzed with the exception of the OFC and ACC, subject age was positively correlated with ICI signal fraction, suggesting that adolescents develop an increased proportion of grey-matter-like tissue in these regions as they age. However, in every reward processing region analyzed, subject age was negatively correlated with ICI signal fraction. This subcortical response to typical development more closely matches previous findings with less specific metrics, though the ICI signal fraction does not necessary correlate with GM volume.

In contrast, increased screen time was associated with a complex series of changes in neural tissue signal fractions in 8 of the ROIs, including 6 regions involved in the control system and 2 involved in reward processing. The only observed association between screen time and ECI was found in the vmPFC, where increased media use had a negative relationship with CSF-like signal fraction. Similar to the association between age and ICA signal fraction, a positive correlation was the only significant relationship observed between screen time and ICA (6 regions). The putamen, amygdala, TPJ, SPL, OFC, and vmPFC were congruent in the associations between screen time and age on ICA signal fraction. These results suggests that increased screen time accelerates the positive trend of ICA development in these regions beyond age-related increases. Significant negative relationships between screen time and ICI signal fraction were observed in 8 of the ROIs. However, these associations were incongruent with the age-related development within cortical control regions (TPJ, IPL, SPL, IFG, OFC, and vmPFC) but congruent with subcortical, reward-processing systems. Thus, these results suggest that screen time has a consistent relationship with microstructure in significant ROIs: accelerating ICA development and decelerating ICI development. However, considering the apparent differences in developmental trajectories between control and reward regions, these results may indicate a fundamental difference in how screen time affects regions involved with the control and reward systems, accelerating development in reward regions and demonstrating mixed responses between signal fractions in control regions.

The contrast between the effects of screen time on tissue microstructure and the effects of age on tissue microstructure characterizes an important relationship between increased screen time and neural development on a cellular level. The relative increase of ICA signal fraction in all significant regions suggests that screen time is associated with a more rapid maturation of axonal innervation in the cortex in these regions. The universal decrease in ICI signal fraction across significant regions is vaguer, but is potentially the result of a decrease in net number of cells in these regions. These results, taken together, suggest that in response to screen time, significantly affected regions of the brain demonstrate a decreasing neural population, increasing axonal myelination, or both simultaneously, indicating that the wiring of the adolescent brain might be maturing more rapidly, possibly at the consequence of the typical neurogenesis observed in the cortex in this age range. However, the cellular responses to increased screen time that underly the relationships observed in this study are relatively ambiguous, as the signal fraction analysis used in this study does not pin changes in tissue microstructure to specific cell types or changes, particularly for ICI signal fraction. Similarly, because this model focuses on relative presence of signal type, whether or not there are effects on the absolute amount of tissue contributing to each signal type cannot be determined.

One possible explanation for the lack of significant FA results in the face of a changing microstructural environment is in the lack of orientation resolution across white matter fibers in this metric. By only modeling unidirectional white matter microstructure in each voxel, the FA model is likely not as sensitive as the 3T-CSD model for this analysis, particularly because of the complex white matter directionality in the cortex. 3T-CSD, however, is able to measure the total signal in a directionless manner. The lack of significant longitudinal effects observed between increased screen time and both GMD and FA, but presence of such results in 3T-CSD metrics highlights both the sensitivity of this technique and importance of multi-dimensional voxel analysis.

While this study found no significant effect of screen time on GMD cross-sectionally, our hypothesis of longitudinal effects, based primarily on the findings of Takeuchi et al. (2018), was not supported by the absence of significant results. Several key differences in the structure of the present study may explain this discrepancy. While the subject age is comparable between studies, the number of subjects and amount of time between MRI visits was different between studies (223 subjects in Takeuchi et al., vs 2666 subjects here, 3 years between scans in Takeuchi et al., vs. 2 years between scans here). Critically, there is also a difference between explanatory variables: Takeuchi et al observing the effects of internet usage, and the present study observing the effects of a host of screen-based activities. While consistent with the current literature, our longitudinal FA results also opposed our hypothesis: only revealing significant relationships between screen time and WM structure in the vmPFC.

There are a host of potential factors influencing the observed relationship between screen time and tissue microstructure, and it is highly likely that the effects of screen time on cellular microstructure in adolescents are related to the contents of screen usage than the medium itself. Possible components that may be driving these changes in cognitive and reward pathways are the over-stimulating and easily gratifying nature of most screen-based activities, which are causing rapid maturation of the reward pathway while inhibiting traditional cognitive development. This explanation supports the negative behavioral and cognitive outcomes that have previously been associated with increased screen usage. In summary, though this study is observational in design, our quantifications of the effects of screen time on neural development in adolescents suggest cellular changes in the brain might be the consequences of habitual media usage. Further research is required to distinguish the specific effects of different screen uses on the adolescent brain, and more rigorously controlled studies manipulating screen time will be necessary to establish causality.

